# Myeloid deletion and therapeutic activation of AMP-activated protein kinase (AMPK) do not alter atherosclerosis in male or female mice

**DOI:** 10.1101/2020.07.15.204487

**Authors:** Nicholas D. LeBlond, Peyman Ghorbani, Conor O’Dwyer, Nia Ambursley, Julia R. C. Nunes, Tyler K.T. Smith, Natasha A. Trzaskalski, Erin E. Mulvihill, Benoit Viollet, Marc Foretz, Morgan D. Fullerton

**Affiliations:** Department of Biochemistry, Microbiology and Immunology, Faculty of Medicine, University of Ottawa, Ottawa, ON, K1H 8M5, Canada; Centre for Infection, Immunity and Inflammation, Ottawa ON K1H 8M5, Canada; Centre for Catalysis Research and Innovation, Ottawa ON K1H 8M5, Canada; University of Ottawa Heart Institute, Ottawa, ON K1Y 4W7, Canada; Université de Paris, Institut Cochin, CNRS, INSERM, F-75014 Paris, France

**Author notes:** To whom correspondences should be addressed: Dr. Morgan Fullerton, Department of Biochemistry, Microbiology and Immunology, Faculty of Medicine, University of Ottawa, 4109A Roger Guindon Hall, 451 Smyth Rd, Ottawa, Ontario, Canada, K1H 8M5.

**Keywords:** AMP-activated protein kinase (AMPK), atherosclerosis, lipids, macrophage, immunometabolism

## Abstract

**Objective:** The dysregulation of myeloid-derived cell metabolism can drive atherosclerosis. AMP-activated protein kinase (AMPK) controls various aspects of macrophage dynamics and lipid homeostasis, which are important during atherogenesis.

**Approach and Results:** We aimed to clarify the role of myeloid-specific AMPK signaling by using LysM-Cre to drive the deletion of both the α1 and α2 catalytic subunits (MacKO), in male and female mice made acutely atherosclerotic by PCSK9-AAV and Western diet-feeding. After 6 weeks of Western diet feeding, half received daily injection of either the AMPK activator, A-769662 or a vehicle control for a further 6 weeks. After 12 weeks, myeloid cell populations were not different between genotype or sex. Similarly, aortic sinus plaque size, lipid staining and necrotic area were not different in male and female MacKO mice compared to their littermate floxed controls. Moreover, therapeutic intervention with A-769662 had no effect. There were no differences in the amount of circulating total cholesterol or triglyceride, and only minor differences in the levels of inflammatory cytokines between groups. Finally, CD68+ area or markers of autophagy showed no effect of either lacking AMPK signaling or systemic AMPK activation.

**Conclusions:** Our data suggest that while defined roles for each catalytic AMPK subunit have been identified, global deletion of myeloid AMPK signaling does not significantly impact atherosclerosis. Moreover, we show that intervention with the first-generation AMPK activator, A-769662, was not able to stem the progression of atherosclerosis.

**Highlights:**

- The deletion of both catalytic subunits of AMPK in myeloid cells has no significant effect on the progression of atherosclerosis in either male or female mice
- Therapeutic delivery of a first-generation AMPK activator (A-769662) for the last 6 weeks of 12-week study had no beneficial effect in either male or female mice
- Studying total AMPK deletion may mask specific effects of each isoform and highlights the need for targeted disruption of AMPK phosphorylation sites via knock-in mutations, rather than the traditional “sledgehammer” knockout approach

## Introduction

Atherosclerosis and its downstream cardiovascular complications continue to represent the leading cause of mortality and morbidity in developed countries. While the importance of cells such as vascular smooth muscle and adaptive immune cells in atherosclerosis has recently been highlighted, myeloid-derived cells of the innate immune system, monocytes and macrophages, are a primary driver of disease initiation and progression^1, 2^. The haematopoietic differentiation of monocytes is crucial for atherogenesis, and it is now appreciated that modulating monocyte pools can have direct effects on atherosclerotic plaque initiation and progression^3^. Moreover, recent lines of evidence have pointed toward the intrinsic metabolic programming of myeloid-derived cells as being a driver of their atherogenic and inflammatory potential^4, 5^.

AMPK is an evolutionarily conserved heterotrimeric serine/threonine kinase that functions to maintain normal metabolic homeostasis by sensing and restoring energy deficits. AMPK acts to limit anabolic and stimulate catabolic programs in the cell. One of the most recognized consequences of AMPK activation is an acute inhibition of both fatty acid and cholesterol synthesis, via inhibiting phosphorylation on acetyl-CoA carboxylase 1 and 2 (ACC1 and 2) and 3-hydroxy-3-methyl-glutaryl-coenzyme A reductase, respectively^6, 7^. The inhibition of ACC results in the reduction of malonyl-CoA, which in addition to regulating lipogenic flux, relieves the inhibition on mitochondrial fatty acid uptake, leading to an increase in β-oxidation^8^. Complementary to this, AMPK, directly and indirectly, stimulates the process of macroautophagy (herein referred to as autophagy), which is critical for processing metabolic substrates, the clearance of damaged or senescent organelles and in the context of atherosclerosis, contributes to the mobilization of stored cholesterol in foam cells and reverse cholesterol transport^9^.

We and others have shown that AMPK signaling in differentiated cultured macrophages can regulate various aspects of fatty acid and cholesterol metabolism, while simultaneously governing broader metabolic and immune programs^10-14^. In the context of atherogenesis, there have been conflicting reports as to the role of myeloid AMPK signaling in the progression of atherosclerosis, though only male mice have been studied^15-17^. Moreover, while systemic delivery of AMPK-activating treatments have been shown to reduce lesion size, the delivery of these treatments was over the entire course of the atherosclerosis progression model^18, 19^ and the contribution of myeloid AMPK was not addressed. Here we report that in an acute model of atherogenesis instigated by AAV-delivery of a gain-of-function PCSK9, myeloid-specific disruption of both the α1 and α2 catalytic subunits of AMPK did not alter plaque size or necrotic core in male or female mice. Further to this, when the direct AMPK activating compound A-769662 was administered to mice in a therapeutic, rather than preventative manner, no protection was observed. These unexpected findings lead us to question the specific roles of each catalytic subunit and whether novel allosteric activators of AMPK would offer therapeutic beneficial effects on atherogenesis.

## Materials and Methods

Materials and Methods are available in the supplementary material online.

## Results

### Deletion of myeloid AMPK signaling does not alter monocyte populations

While AMPKα1 is the predominant isoform expressed in haematopoietic cells^12, 20^, myeloid deficiency of either AMPKα1 or AMPKα2 has been shown to alter atherosclerosis^15-17^. Moreover, these studies crossed their respective AMPK-deficient mouse models onto the atherogenic *LDLr*- or *ApoE*-deficient background. To completely disrupt AMPK signaling, we generated mice lacking all AMPK signaling in myeloid cells (AMPKα1 and AMPKα2 deletion driven by LysM expression of Cre recombinase), which was confirmed by assessing AMPK-specific signaling to its downstream target ACC in elicited peritoneal macrophages from floxed littermate controls (wild-type; WT) and MacKO mice. As expected, basal AMPK signaling to ACC was almost completely absent in MacKO cells, with no effect of the AMPK activator, A-769662 (Supplemental Figure S1). As signaling to ACC has long been shown to be AMPK-specific^21^, the residual signal is likely due to the presence of non-myeloid cells. Importantly, basal AMPK signaling in the liver was unaffected (Supplemental Figure S1).

To circumvent genetic mouse models of atherosclerosis, we used a single intravenous injection of a well-characterized, gain-of-function (D377Y) mouse PCSK9-AAV followed by Western diet (WD) feeding to induce rapid hypercholesterolemia to drive atherogenesis for 12 weeks (Supplemental Figure S2)^22^. Monocyte-derived macrophages are a key cell type in atherosclerosis, and myeloid cell maturation and differentiation, which begins in the bone marrow via myelopoiesis can be a driver of atherosclerosis. Moreover, myelopoiesis is augmented by WD-induced hypercholesterolemia in both mice and humans^23^. In a preliminary cohort of male and female WT and MacKO mice, we first aimed to determine if there were any basal genotype differences in myeloid cell populations. Using Ly6C, CD115 and Ly6G as myeloid cell markers, we observed no statistical differences in circulating, splenic or bone marrow populations, either between genotypes or between males and females (Figure 1A and Supplementary Figure S3). Moreover, the presence or absence of myeloid AMPK did not skew the levels of Ly6C^hi^ or Ly6C^lo^ populations, which are recognized as monocytes prone (Ly6C^hi^) or resistant (Ly6C^lo^) to effector or patrolling functions, respectively^2^ (Figure 1B and C). To investigate myeloid differentiation, we determined that bone marrow Lin^−^Sca-1^+^c-Kit^+^ cells, as well as bone marrow and splenic populations of multipotent progenitors, common myeloid progenitors, granulocyte-macrophage progenitors and megakaryocyte-erythroid progenitors were also unaltered (data not shown).

**Figure 1.**
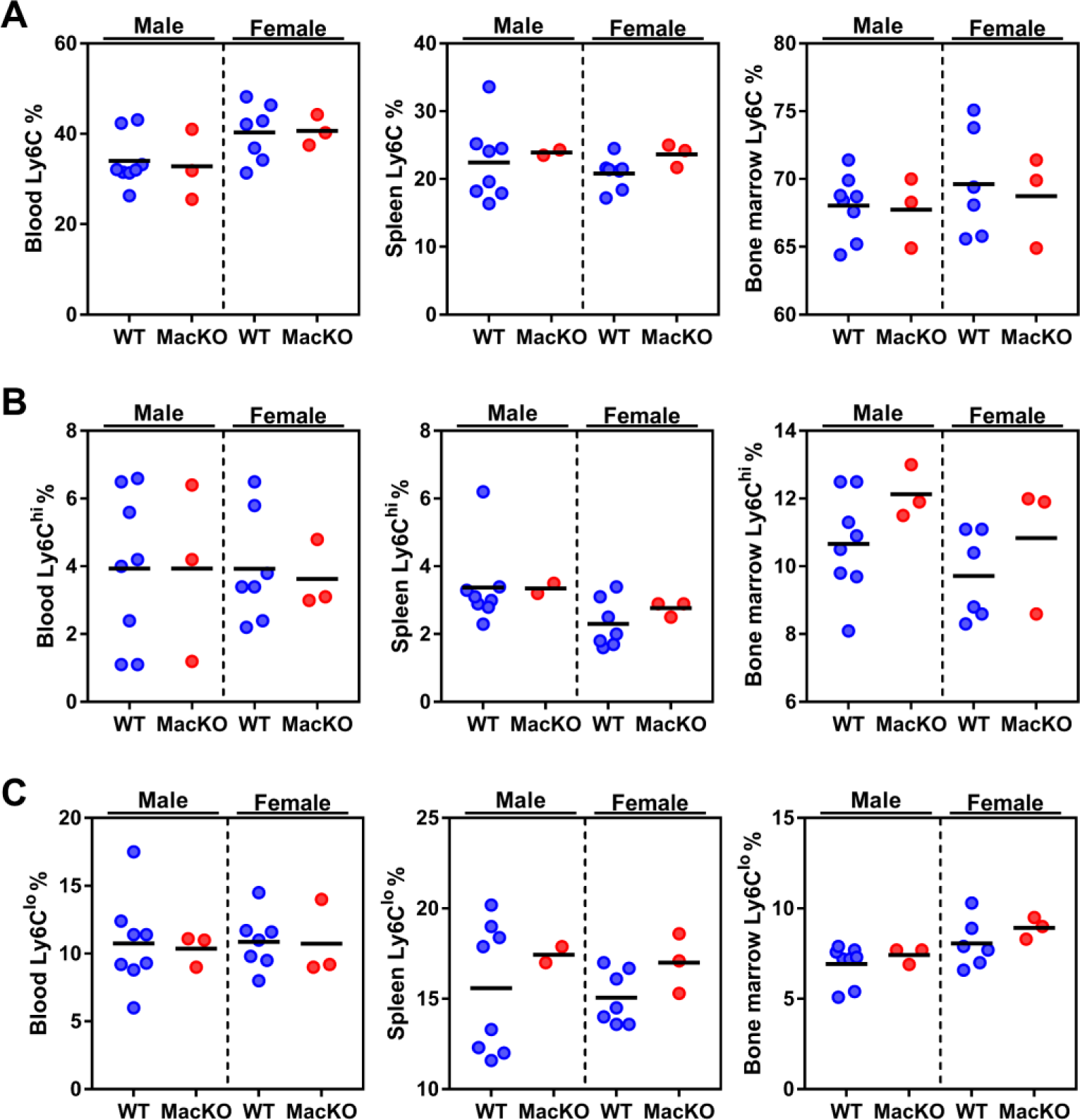
Myeloid AMPK does not alter myelopoiesis. WT and MacKO male and female mice were injected with the PCSK9-AAV and fed a WD for 12 weeks. The expression of (A) total Ly6C, (B) Ly6C^hi^ and (C) Ly6C^lo^ populations from the blood, spleen, and bone marrow quantified. Each data point represents the mean value from one animal (n = 3-8/group).

### Myeloid AMPK signaling has minimal impact on the atherosclerotic plaque

Male and female mice of both genotypes gained weight as expected when fed a WD. After 6 weeks, each group was divided and received daily injections with either vehicle control or the first-generation AMPK activator A-769662 (30 mg/kg) for the last half of the 12-week intervention. Both male and female mice, independent of myeloid AMPK signaling experienced a reduction in body weight, as has been previously documented^24^ (Supplemental Figure S4). At the completion of the 12-week study, the atherosclerotic lesion area was quantified from the aortic sinus. In male and female mice, there were no differences in lesion size between genotypes. In addition, treatment with A-769662 as an intervening therapy had no significant effect on plaque size (Figure 2A). When plaque sections were stained with Oil Red O to quantify neutral lipid, there were no significant differences between groups (Figure 2B). Consistently, the amount of necrotic area was also not changed between by the presence of myeloid AMPK, treatment with A-769662 or between sex. (Figure 2C and Supplemental Figure S5). These data suggest that in the early stages of PCSK9-induced atherosclerosis, myeloid AMPK signaling does not regulate total lesion area, lipid content or necrotic area.

**Figure 2.**
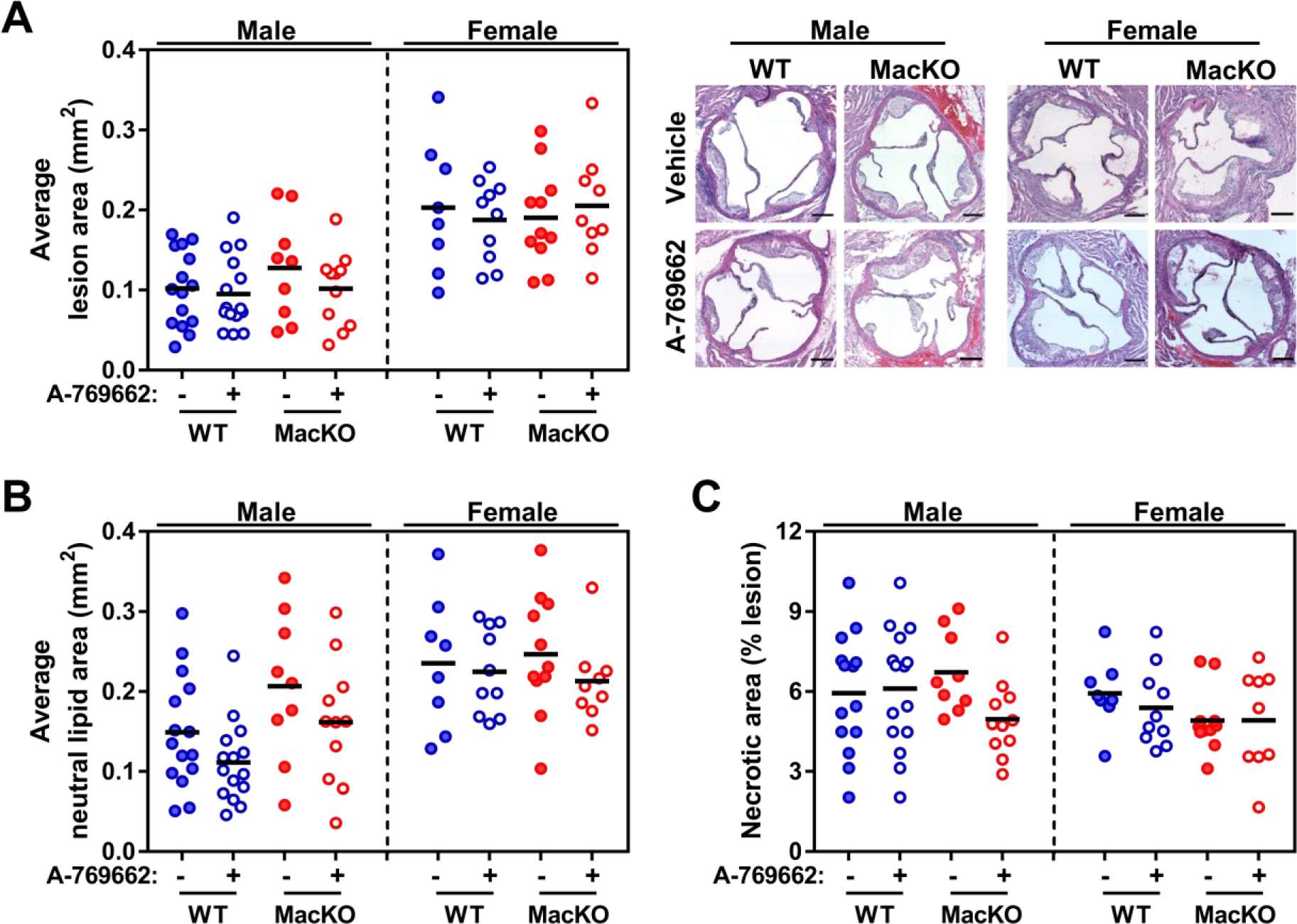
Myeloid AMPK signaling does not alter the progression of atherosclerosis in male and female mice. WT and MacKO male and female mice were injected with the PCSK9-AAV and fed a WD for 12 weeks, with half of each group receiving either 30 mg/kg A-769662 or vehicle control for the final 6 weeks. (A) Average lesion area quantified within the aortic sinus (representative H&E-stained images are included). (B) Quantification of the average oil red O containing areas within the aortic sinus. (C) Lesion necrotic area expressed as percent lesion area. (A-C) Selection and quantifications were performed with Image J software of (A and C) H and E and (B) Oil Red O stained slides as listed in the materials and methods. Each data point represents the mean value from one animal (n = 7-16/group).

### Myeloid AMPK and systemic activation do not affect total circulating lipid levels

The liver is a significant regulator of whole-body lipid metabolism and can dictate levels of lipoprotein-associated cholesterol and triglyceride in the circulation. While disruption of AMPK was restricted to myeloid cells, there is the potential that LysM-mediated deletion would occur in liver-resident macrophages (Kupffer cells), which could affect circulating lipid levels^25^. Moreover, our experimental design made use of the systemic delivery of A-769662, which has well known effects on lipid metabolism^21, 24^. Independent of genotype, treatment or sex, there were no differences in the levels of circulating total cholesterol or triglyceride (Figure 3A and B).

**Figure 3.**
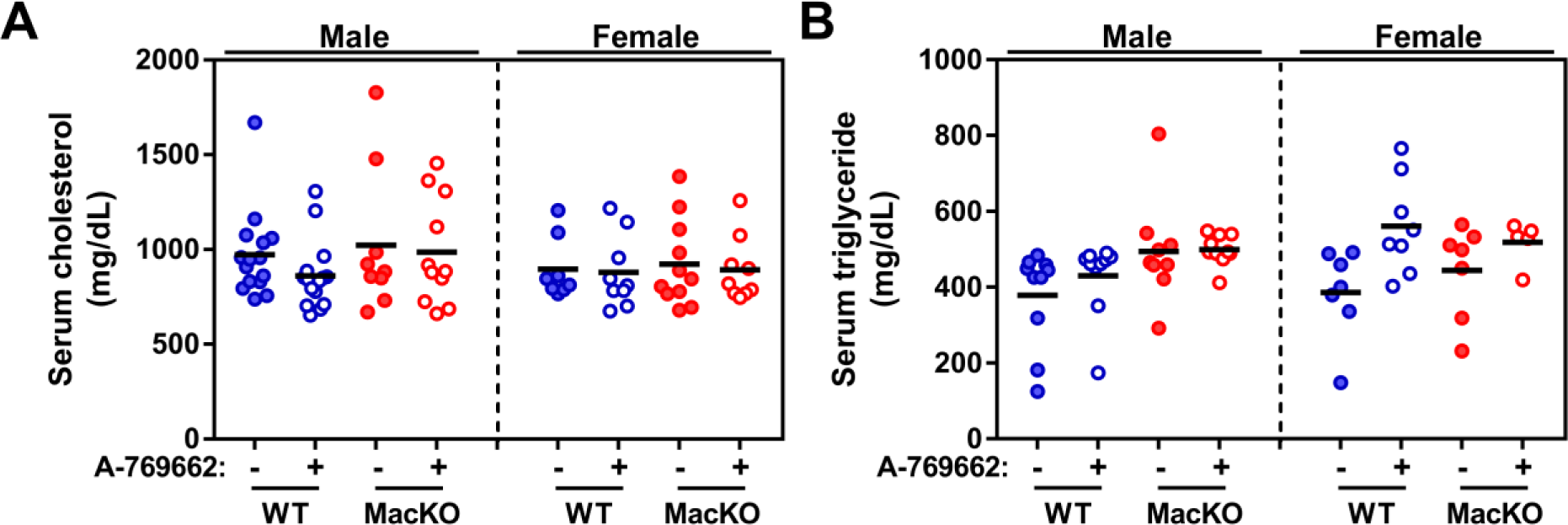
Myeloid AMPK signaling does not change total levels of circulating lipids. Following WD-feeding, serum was collected to assess (A) total circulating cholesterol and (B) total circulating triglycerides. Each data point represents the value from one animal (n = 5-16/group).

### Myeloid AMPK does not regulate systemic inflammation during atherosclerosis

Metabolism and inflammation are recognized as important drivers of atherogenesis. Given the known role for AMPK in modulating both macrophage metabolism and immune programs, we next assessed the systemic (circulating) levels of inflammatory cytokines (IL-10, IL-12p70, IL-6, KC, TNFα, IL-1β, IFNγ, IL-2, IL-4 and IL-5) (Figure 4 and Supplemental Figure S6). In male mice, at the time of tissue harvest and blood collection, circulating levels of IL-12p70 and IL-4 were significantly lower in MacKO compared to control mice. When comparing vehicle and A-769662 treatment, IL-12p70 levels decreased in WT-treated but increased in MacKO-treated mice (Figure 3A-D). In addition, levels of KC (mouse IL-8), were significantly increased in A-769662-treated WT mice, but lower in MacKO mice treated with the AMPK activator. While levels of other cytokines went unchanged, IL-12p70, IL-4 and IL-10 were all augmented by A-769662 treatment in mice that were deficient for myeloid AMPK, suggesting non-AMPK or non-myeloid mechanisms. In female animals, only IL-12p70 differed significantly, such that levels in MacKO mice were lower, but that also in WT mice, A-769662 treatment also caused a reduction in circulating levels.

**Figure 4.**
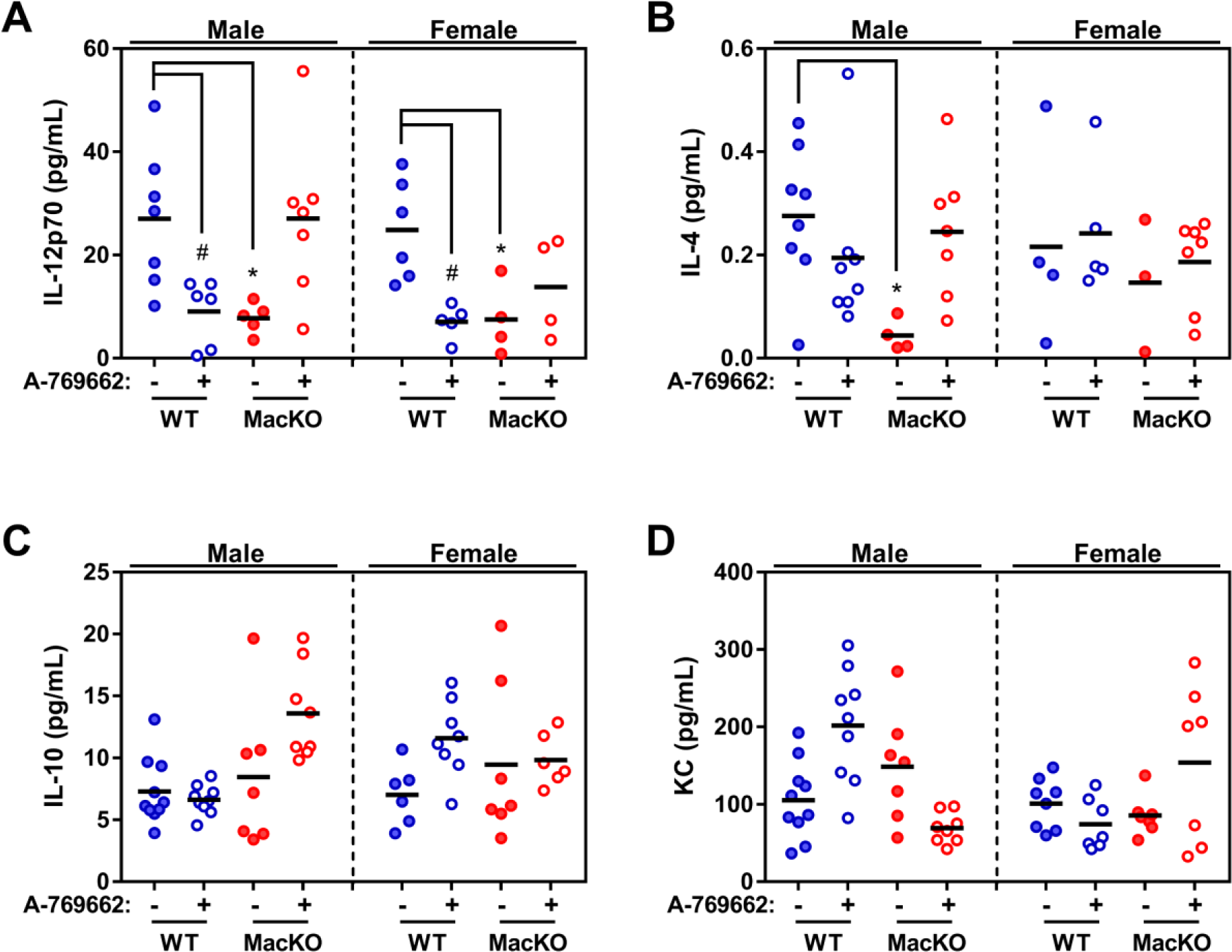
Myeloid AMPK signaling slightly alters systemic inflammation during the progression of atherosclerosis. Following WD-feeding, serum was collected to assess circulating inflammatory cytokines (A) IL-12p70, (B) IL-4, (C) IL-10, and (D) KC. Each data point represents the value from one animal (n = 4-10/group), where * represents p<0.05 between genotypes and # is p<0.05 between treatments as calculated by a 2way ANOVA.

Previous work has demonstrated that preventative AMPK activation (i.e. AMPK activator treatment at the start of dietary initiation of atherosclerosis) was associated with decreased levels of the circulating chemokine monocyte chemoattractant protein 1 (MCP1)^18, 26^. Consistent with a lack of difference in total plaque area, circulating MCP1 was not different between any of the groups (Figure S7).

### Myeloid AMPK signaling does not alter the amount of CD68+ cells

Deletion of myeloid AMPK on an *ApoE*-deficient^16^ or *LDLr*-deficient^17^ background resulted in less and more markers of macrophage accumulation, respectively. To interrogate this in our model, we used CD68 as a general marker of macrophage-like cells within atherosclerotic lesions, since it is now well established that vascular smooth muscle cells can adopt CD68 expression as atherogenesis progresses^27^. Similar to results above, we observed no difference in aortic lesion area between male WT and MacKO, we detected no difference in the amount of lesion CD68+ expression between groups, regardless of genotype or treatment when normalized to total area (Figure 5A and B).

**Figure 5.**
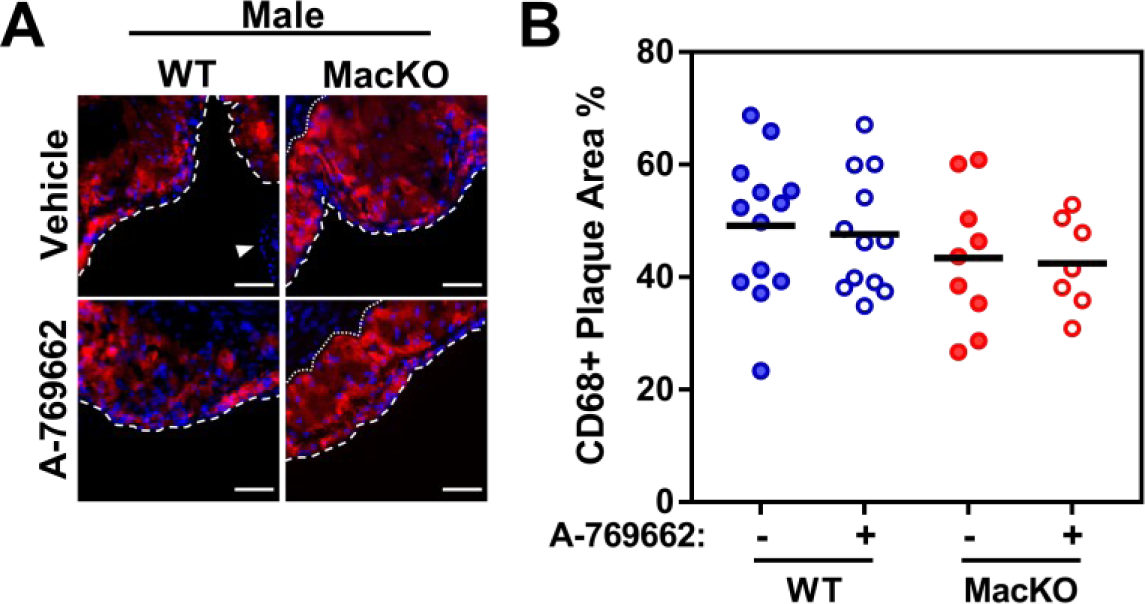
Myeloid AMPK signaling does not regulate the proportion of CD68-positive cells within the lesion but is implicated in total CD68 expression within male mice. (A) Representative images for immunofluorescent labeled lesions of male mice (CD68; Red, DAPI; Blue, where the luminal border of the lesion is denoted by white dashed and dotted line). (B) Quantification of CD68+ area as a percent of the total lesion area in lesions of male mice. Scale bar represents 50 µm. Each data point represents the mean value from one animal (n = 7-13/group).

### Myeloid AMPK signaling does not alter markers of autophagy

We next aimed to determine if the disruption or activation of AMPK signaling in myeloid cells influenced markers of autophagy within the plaque environment. We began by probing for the expression of autophagy markers Beclin-1 and microtubule-associated proteins 1A/1B light chain 3B (LC3II/I) in whole aortic lysates of male WT and MacKO mice; however, no basal genotype differences were observed (Figure 6A). We next stained the atherosclerotic lesion for p62 (also known as SQSTM1), an essential chaperone protein that is processed in an autophagic-dependent matter and serves as a marker of defective autophagy^28^. The average level of plaque-associated p62 was unaffected by the presence or absence of myeloid AMPK, and treatment with A-769662 had no effect on the levels of p62 staining within lesions (Figure 6B and C) in male mice. We also observed that the localization of p62 staining within the plaque, which has been linked to its potential sequestration along with its cargo to autophagosomes, was also not regulated by genotype or treatment.

**Figure 6.**
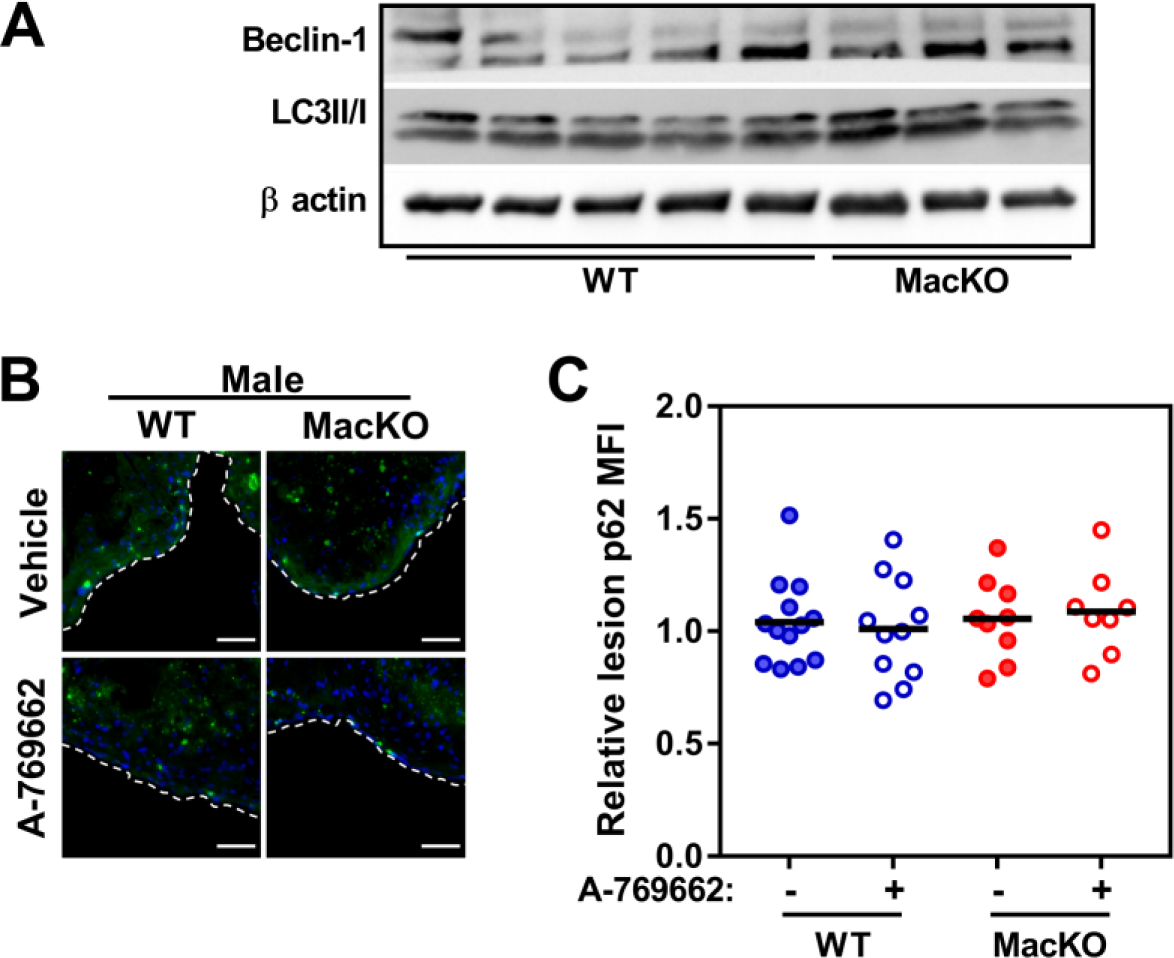
Myeloid AMPK signaling does not influence markers of lesion autophagy. (A) Whole aortic protein lysate from WT and MacKO (male) mice was probed for the expression of Beclin-1 and LC3II/1. (B) Close up representative image for immunofluorescent labeled lesions of male mice (p62; Green, DAPI; Blue, where the luminal border of the lesion is denoted by a white dashed line). (C) Quantification of the relative mean fluorescent intensity for p62 within the lesions of male mice. Scale bar represents 50 µm. Each data point represents the mean value from one animal (n = 8-13/group).

## Discussion

The progression of atherosclerosis is causative in the development of cardiovascular disease. While mechanistic understanding has translated to effective frontline therapies against cardiovascular disease, incidence and societal burden remain high. Numerous pathways and cell types are engaged at the onset of atherosclerosis; however, among the complexity, metabolic and cell differentiation pathways of myeloid cells are critical^2^. Since AMPK is known to regulate multiple immunometabolic programs^8, 29^, we sought to address the importance of AMPK signaling in myeloid cells during the progression of atherosclerosis. Moreover, building on evidence that chronic, preventative treatment of WD-fed *ApoE*-deficient mice with AMPK activators (including A-769662) protected against lesion development^18, 19^, we asked whether therapeutic intervention with systemic AMPK activation protects against atherogenesis in a myeloid AMPK-dependent manner.

To avoid lengthy breeding schemes and the constitutive nature of *Ldlr*^*-/-*^ and *ApoE*^*-/-*^ mouse models, we administered a validated gain-of-function (D377Y) PCSK9-AAV, which was driven by a liver-specific promoter^22^. While we aimed to provide clarity as to the role of myeloid AMPK in atherosclerosis, in our model, there were no statistical differences in plaque, lipid or necrotic area between 1) mice that had or did not have myeloid AMPK signaling, or 2) mice that were treated with or without the direct AMPK activator A-769662, in male and female mice. However, we observed that in male mice only, myeloid AMPK signaling showed a trend toward restraining the lipid content of atherosclerotic plaque in an AMPK-dependent manner (Figure 2B). This finding, though not statistically significant, is consistent with previous observations showing a protective role for AMPK signaling when the AMPKα1 subunit was deleted using LysM-Cre on an *Ldlr*-deficient background^17^. Similarly, we found that intervention with an AMPK activator tended to stem plaque lipid accumulation and that myeloid AMPK signaling seemed to be required.

There is no clear answer to the question as to whether AMPK and more specifically, myeloid AMPK plays a beneficial or detrimental role in atherogenesis. When myeloid AMPKα1 deletion was driven by LysM expression on a *Ldlr*-deficient background, these mice had more atherosclerotic lesion and plaque area, higher circulating lipids, more Ly6C^hi^ monocytes and more pro-inflammatory markers in the aorta^17^. Opposing this, myeloid AMPKα1 or AMPKα2 deletion via LysM-mediated Cre expression on an *ApoE*-deficient background resulted in smaller atherosclerotic lesions. This was attributed to an AMPKα1-dependent effect on regulating monocyte-to-macrophage differentiation and autophagy^16^ or an AMPKα2-dependent effect on DNA methylation and altered expression of matrix metalloproteinases^15^, respectively. The phenotype of the myeloid AMPKα2-deficient mice is interesting given that the predominant isoform in myeloid cells was shown to be AMPKα1^12, 30^. Given this, it might be expected that a compensatory up-regulation of the AMPKα2 subunit might occur during AMPKα1-deficiency; however, the opposite is not as intuitive.

Rather than deal with isoform-specific contributions, we chose to delete both catalytic subunits to best characterize the significance of the sum of myeloid AMPK signaling. One potential limitation to all studies that have used LysM-Cre as a driver of myeloid-specific disruption is the chronic and constitutive nature of the targeted deletion. However, even when this is considered, the largest discrepancy between studies that show a protective and those that show a detrimental role for myeloid AMPK is genetic background (LDLr vs. ApoE). Atherosclerosis is lessened in response to myeloid AMPKα1 and AMPKα2 deletion only when on an *ApoE*-deficient background^15, 16^. Moreover, AMPKα1/ApoE mice had no difference in serum cholesterol and AMPKα2/ApoE mice had higher total cholesterol levels compared to floxed/ApoE control mice, suggesting that these effects were independent of circulating cholesterol. Contrary to this, circulating cholesterol levels in AMPKα1/LDLr mice were positively correlated with lesion size and were increased when myeloid AMPKα1 was absent^17^. We did not observe any differences in total levels of circulating lipids, which was consistent with the similarity in total lesion area in all groups. Yet lesion lipid accumulation in male myeloid AMPK-deficient mice trended toward being higher in our PCSK9-induced model (Figure 2B).

There remains the potential that regulatory differences in AMPK signaling exist in myeloid cells that have or do not have LDLr or ApoE. Aldehyde dehydrogenase 2 (ALDH2) disruption was atheroprotective on an *ApoE*-deficient background but atherogenic on a *Ldlr*-deficient background^31^. The presence of the LDLr blocks AMPK from phosphorylating ALDH2, which leads to histone deacetylase 3 (HDAC3)-mediated down-regulation of lysosomal programs and increased foam cell formation^31^. Two considerations are important when applying this to our current findings. First, myeloid deletion of AMPK signaling would remove the negative signaling to HDAC3 and serve to enhance programs that lower atherogenesis. Secondly, in our PCSK9-induced model, whether circulating PCSK9 has LDLr-suppressive effects in non-hepatic tissues (specifically circulating monocytes and plaque macrophages) remains unknown, making any interpretation of myeloid AMPK signaling working via LDLr premature.

AMPK signaling has been shown to induce autophagy directly via activating phosphorylation of Unc-51 like autophagy activating kinase 1, Beclin-1 and VPS34, and indirectly via inhibition of the mechanistic target of rapamycin complex 1, which itself is regulated indirectly by AMPK-mediated phosphorylation of tuberous sclerosis complex 2 and/or regulatory-associated protein of mTOR^32^. It is now well established that autophagy is athero-protective due to its clearance of cellular debris, sequestration of defective organelles and liberation of free cholesterol for the purpose of cholesterol efflux^33^. To assess one marker of lesion autophagy, we stained lesions for p62, the accumulation of which is associated with defective autophagy. Despite initial predictions, there were no genotype or treatment effects on the expression or localization p62 within the developed lesions. With a trending decrease in lesion lipid content, it remains entirely plausible that the effects of AMPK signaling are linked to its regulation of autophagy; however, this was not captured by p62 staining. In isolated macrophages, we and others have demonstrated that in addition to an acute regulation of autophagy induction, AMPK exerts a level of transcriptional regulation via direct and indirect modulation of transcription factor EB and lysosomal programs^34-36^.

Previous reports show that genetic deletion of myeloid AMPK primes them toward a more pro-inflammatory phenotype, elevating proportions of Ly6C^hi^ to Ly6C^lo^ monocytes in the circulation or in the peritoneum in response to thioglycollate^17^. In keeping with this, mice treated systemically and chronically with A-769662 had a similar response and lower Ly6C^hi^ cells^18^. In the present study, with all AMPK signaling disrupted, we observed absolutely no effect on circulating, splenic or bone marrow populations of myeloid or precursor cells. This was consistent with the lack of difference regarding plaque size but differ with results from *ApoE* and *Ldlr*-deficient models^15-17^. As a correlate of immune infiltration into the plaque, we used CD68 as a marker of macrophages and macrophage-like cells, although CD68 is also known to be expressed on dendritic cells, neutrophils and smooth muscle cells as well^27^. Unexpectedly, we observed no differences in the apparent proportion of lesion associated CD68+ cells. While chronic A-769662 treatment in *ApoE*-deficient mice resulted in lower Ly6C^hi^ monocytes in conjunction with decreased C-C chemokine receptor type 2 (*Ccr2*) expression, this was recently expanded by the observation that in response to acute fasting or acute (4 h) AMPK activation with A-769662, Ly6C^hi^ monocytes are dramatically reduced due to lower levels of MCP1, the ligand for CCR2^26^. In our model of complete myeloid AMPK-deletion, we see that circulating MCP1 is not altered. Importantly, while the effects of acute fasting and AMPK activation were attributed to a hepatic AMPK signaling, we did not observe any differences in MCP1 levels following systemic AMPK activation.

There have been an impressive number of studies that have aimed to assess the contribution of AMPK toward atherosclerosis. Whole-body^16, 37, 38^, vascular smooth muscle^39, 40^, endothelial^39, 41^, and myeloid-specific models on either *ApoE*- or *Ldlr*-deficient backgrounds^15-17^ have been used in conjunction with either AMPKα1 or AMPKα2 knockout models. Despite this, to our knowledge, female mice have rarely (if ever) been assessed. Our results suggest that while there are no significant differences between male and female MacKO mice and WT littermates, there is a potential sex-specific trend whereby AMPK signaling may act to lower lesion lipid content in male, but not female mice. Sex-specific responses will continue to be an essential consideration in any future work. Moreover, a potential interpretation could be that the deletion of both catalytic subunits of AMPK, the effects (beneficial or detrimental) observed in models of single AMPK deletion may be lost. This point highlights the inherent limitations in knockout models of important metabolic regulators and the need for more pointed models to tease out the importance of specific signaling nodes.

We feel it is important to address some of the limitations and caveats of our study. 1) We began by addressing how myeloid AMPK deficiency may alter immune populations; however, we did not assess mice following the intervention with A-769662. While this may have illuminated differences in monocyte differentiation, there were no statistical differences in atherosclerosis. 2) Also, we chose to inject our WT and MacKO mice with a gain-of-function PCSK9 AAV to induce atherosclerosis (2×10^10^ viral genomes per mouse), which at this concentration when coupled with the WD-feeding, resulted in levels of circulating cholesterol and plaque sizes that were lower than what is typically observed in *LDLr*-deficient mice^22^. Though 12 weeks of atherosclerosis progression should be sufficient for genotype differences to emerge, it may be possible that differences could have been observed earlier or later. Moreover, although treatment with A-769662 daily during atherogenesis was shown to be protective, intervention at the same dose was mainly ineffective and suggests that AMPK-activation may be necessary from the start of atherogenesis to provide a therapeutic benefit. However, this does not rule out the possibility that second-generation allosteric AMPK activators (PF-739, PF-249 or MK-8722), which are more potent and specific AMPK activators, will not show a therapeutic benefit during an intervention. This will be an important consideration for future work.

Our study is the first to question the role of both AMPKα1 and AMPKα2 in myeloid cells and suggests that the presence and activation of myeloid AMPK signaling does not impact atherosclerotic plaque size in the aortic root, independent of sex. Importantly, there were no changes in myeloid cell numbers, circulating lipid levels or systemic inflammation. While we have taken yet another approach to assessing the contribution of AMPK to atherogenesis (PCSK9-induced atherosclerosis and deletion of both myeloid AMPKα1 and AMPKα2), future work should concentrate on dissecting the specific molecular and metabolic pathways by which this important regulator may act. Moving forward, targeted phosphorylation knock-in mouse models of known AMPK substrates will be the only way to specifically interrogate the mechanism(s) by which AMPK-regulated pathways like autophagy and lipid metabolism affect atherosclerosis and other cardiometabolic diseases.

## Acknowledgements

We would like to thank Xiaoling Zhao for her assistance with histology and Dr Katey Rayner for her guidance with aortic sectioning.

## Abbreviations

ACC1/2: Acetyl-CoA carboxylase 1 and 2
ALDH2: Aldehyde dehydrogenase 2
AMPK: AMP-activated protein kinase
CCR2: C-C chemokine receptor type 2
HDAC3: Histone deacetylase 3
LC3 I/II: Microtubule-associated proteins 1A/1B light chain 3B
MacKO: AMPKα1/α2 flox-LysM Cre+
MCP1: Monocyte chemoattractant protein 1
p62: Ubiquitin-binding protein p62 or Sequestosome-1
WD: Western diet
WT: Wild-type; AMPKα1/α2 flox-LysM Cre-

