## Supplementary material for "Myeloid deletion and therapeutic activation of AMP-activated protein kinase (AMPK) do not alter atherosclerosis in male or female mice"

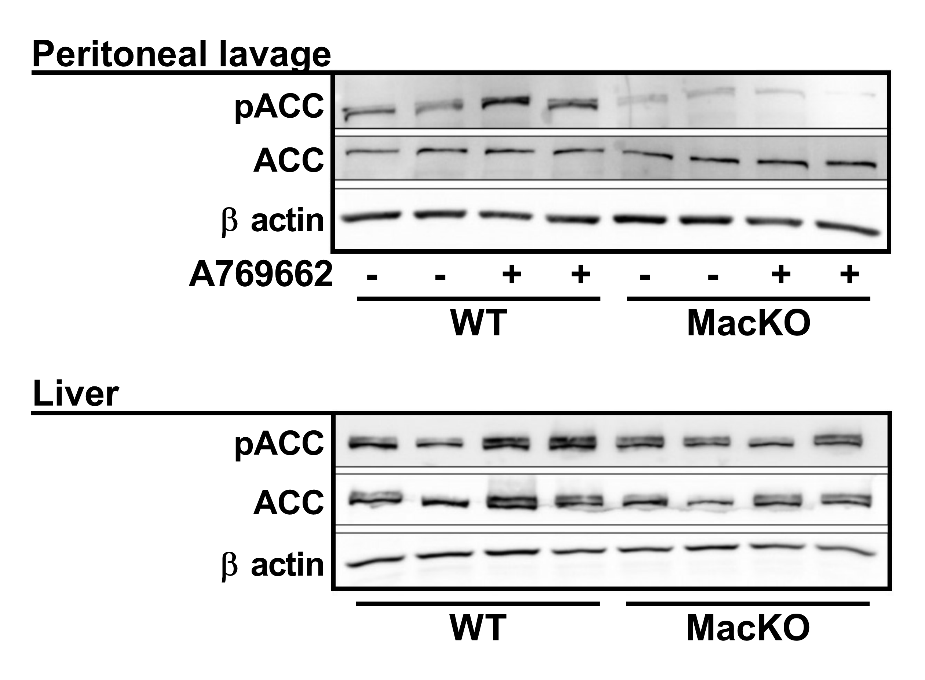

**Supplementary Figure S1.** Validation of myeloid AMPK-deficiency. Representative immunoblot depicting AMPK-specific signaling to ACC at S79 in peritoneal isolated cells (top) and liver (bottom). All mice received an I.P. injection of 3% thioglycolate four days prior to the isolation of cells from the peritoneal cavity and liver tissue. Peritoneal cells were harvested and incubated ± 100 µM A-769662 for 5 hours, whereas the liver was removed and probed without treatment. Total and phosphorylated ACC were assessed from duplicate gels.

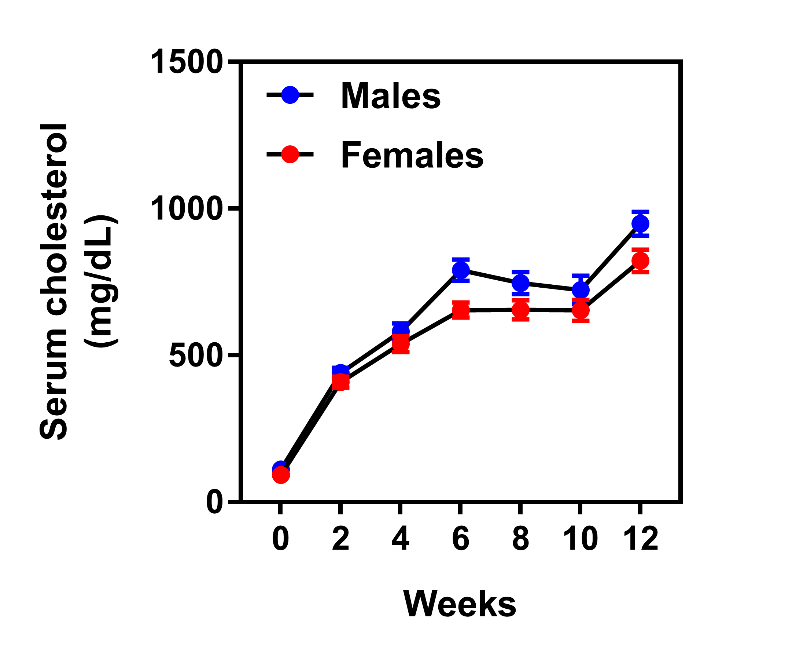

**Supplementary Figure S2.** WD-fed mice infected with PCSK9-AAV become hypercholesterolemic. Total blood cholesterol was measured weekly and shown as a pooled, representative group of male and female mice. Each data point represents the mean value from one animal ± SEM (n = 9-16/group).

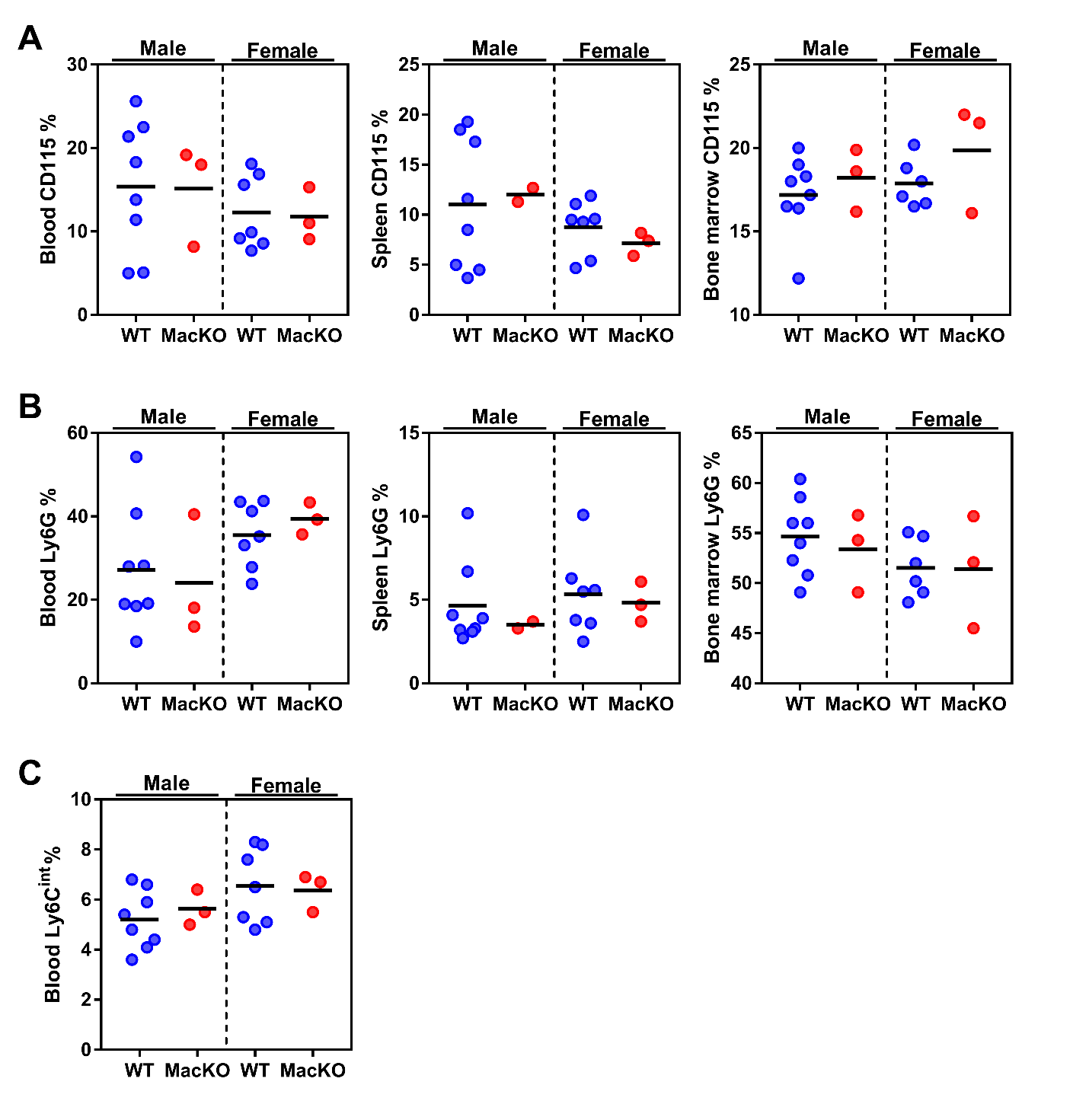

**Supplementary Figure S3.** Myeloid AMPK does not alter CD115 and Ly6G expressing cell populations. WT and MacKO male and female mice were injected with the PCSK9-AAV and fed a WD for 12 weeks. Cells expressing (A) CD115 and (B) Ly6G were quantified from the blood, spleen, and bone marrow. (C) Percentage of Ly6C^int^ expressing cells in circulation. Each data point represents the mean value from one animal (n = 2-8/group; a MacKO sample from the spleen was removed due to improper staining).

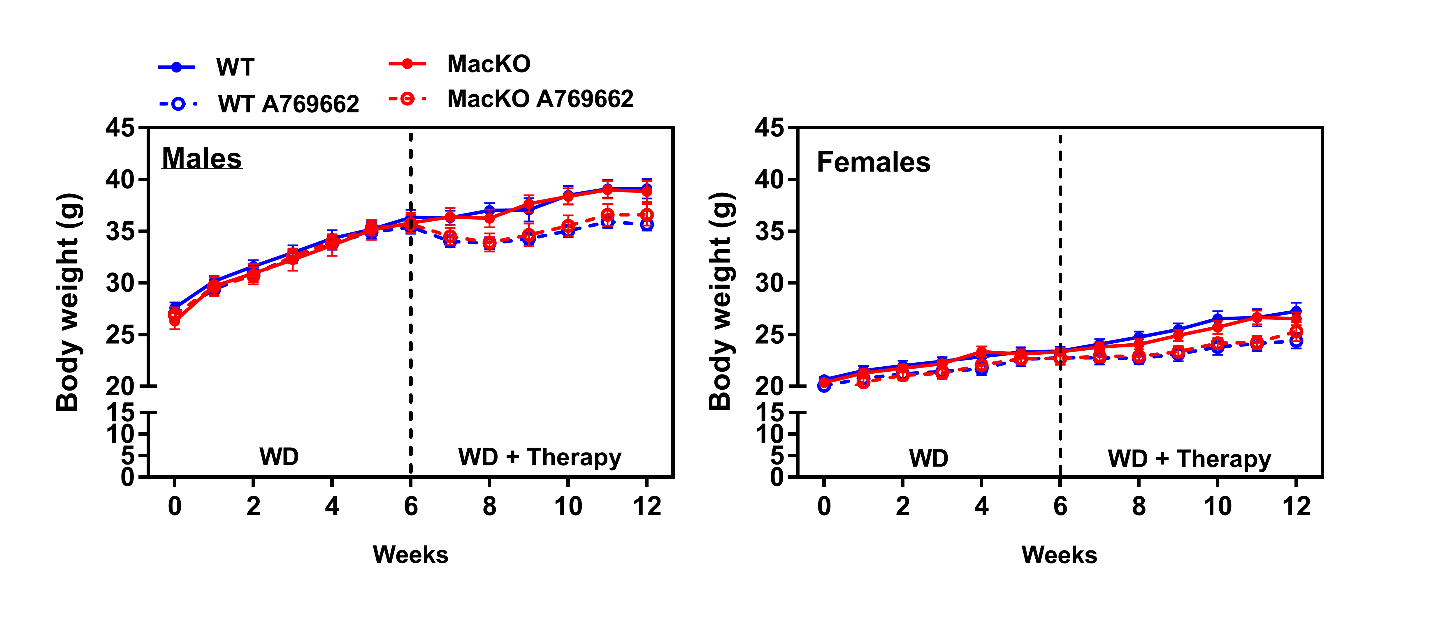

**Supplementary Figure S4.** Weekly mouse weights were monitored post-PCSK9-AAV injection for (A) male and (B) female WD-fed mice. At 6-weeks of HF-feeding mice were placed into groups and treated daily with either 30 mg/kg A-769662 or PBS control (I.P). Each data point represents the mean value from one animal ± SEM (n = 9-16/group).

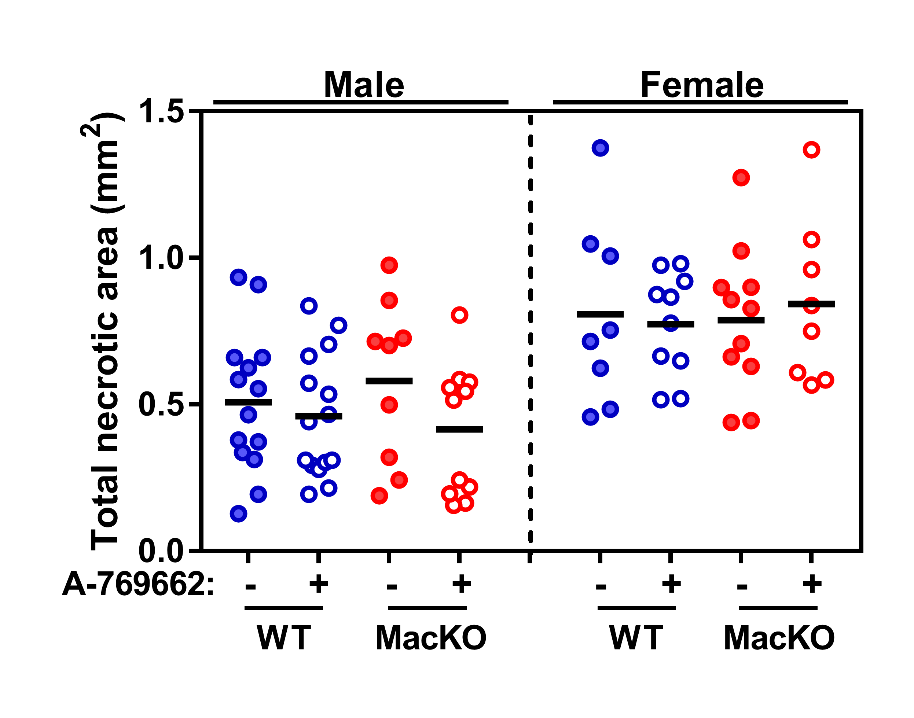

**Supplementary Figure S5**. The total necrotic area (denoted as white space absent of cellularity) was assessed in the lesions of male and female mice. Each data point represents the mean value from one animal (n = 8-15/group).

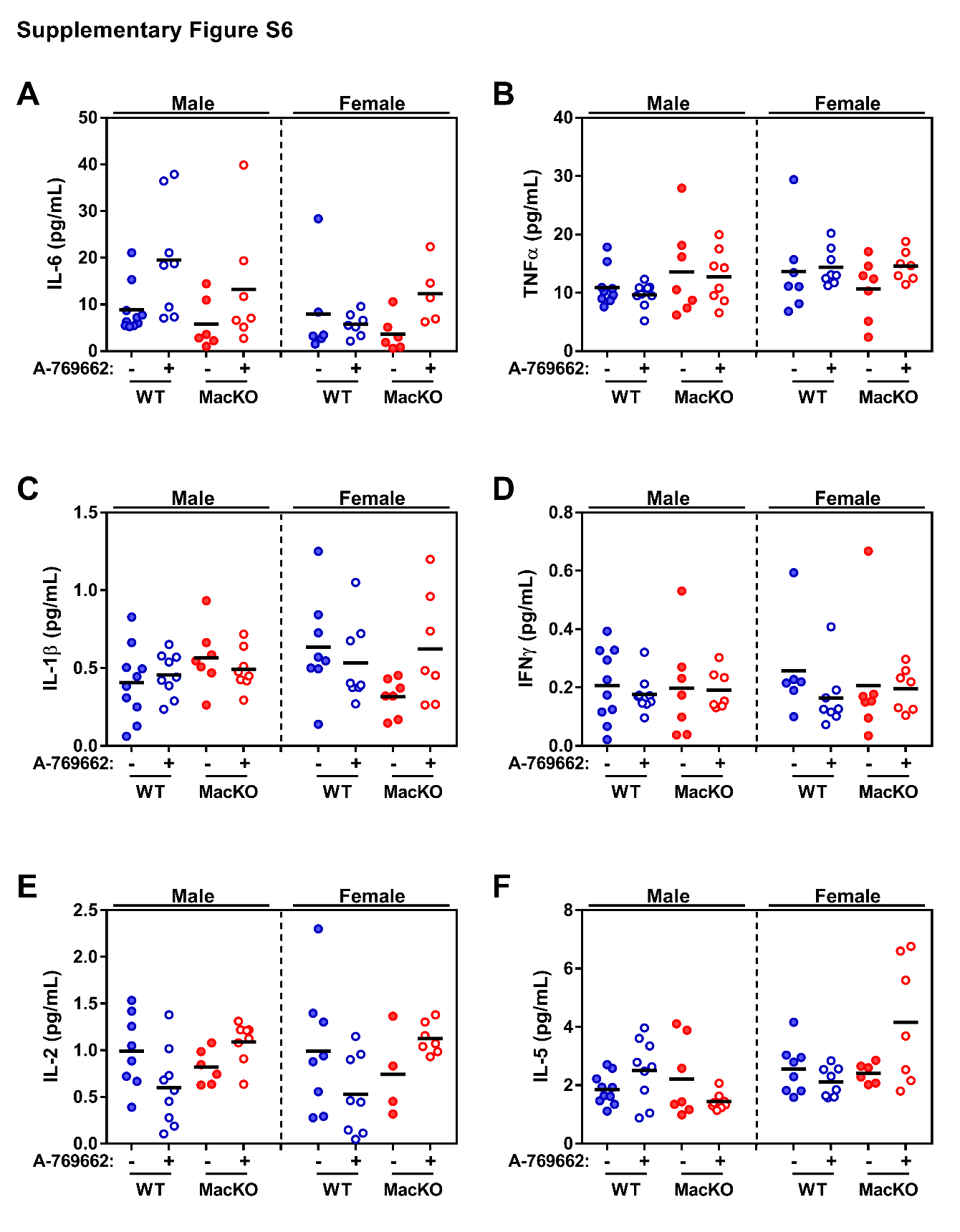

**Supplementary Figure S6.** Myeloid AMPK signaling does not alter systemic cytokine levels. Quantification of circulating inflammatory cytokines (A) IL-6, (B) TNFα, (C) IL-1β, (D) IFNγ, (E) IL-2, and (F) IL-5 was performed on endpoint serum samples obtained during tissue harvest. Each data point represents the value from one animal (n = 4-10/group; certain samples were below the detectable levels for select cytokines).

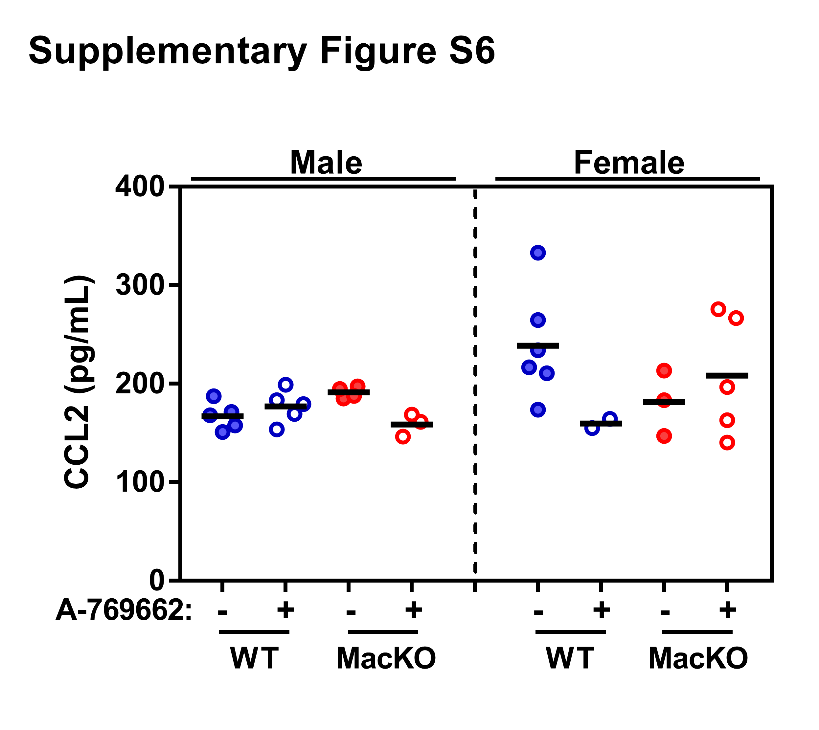

**Supplementary Figure S7.** Circulating levels of MCP1 measured from endpoint serum obtained during tissue harvest. Each data point represents the value from one animal (n = 2-6/group; certain samples were below the detectable levels for select cytokines).

**Materials and Methods:**

**Animal Studies**:

All experiments conducted were in accordance with the Canadian Council of Animal Care and approved by the Animal Care Committee at the University of Ottawa. All mice were housed in ventilated cages at ~23 °C and maintained on a 12/12 h light-dark cycle with *ad libitum* access to a standard rodent chow (44.2% carbohydrates, 6.2% fat, and 18.6% crude protein; diet T.2018, Harlan Teklad). The generation of AMPKα1^flox^ and AMPKα2^flox^ mice has been previously described^20, 21^. AMPKα1^flox^ and AMPKα2^flox^ mice were crossed to obtain double AMPKα1^flox^/α2^/flox^ animals and then crossed with lysozyme Cre (LysM-Cre) hemizygous mice^22^. Littermate animals were used for these studies by only breeding AMPKα1^flox^/α2^/flox^ to AMPKα1^flox^/α2^/flox^ LysM-Cre positive (herein referred to as WT and MacKO). All mice described here are on a C57Bl/6J background. At 10 weeks of age, mice were injected intravenously with 2.5 x10^10^ genome copies of a mouse PCSK9-AAV (D377Y). This plasmid was obtained from Addgene (plasmid #58376, which was a kindly deposited by Dr. Jacob Bentzon^23^), and packaged in an AAV8 by the Penn Vector Core at the University of Pennsylvania. Immediately following viral infection, mice were placed on a Western diet (40% kcal from fat, 43% kcal from carbohydrate, 17% kcal from protein, with 0.2% cholesterol – Research Diets D12079) to induce hypercholesterolemia. After 6 weeks, male and female WT and MacKO mice were assigned to groups that received daily intraperitoneal injections of either 30 mg/kg A-769662 or vehicle control (5% DMSO in PBS) for a further 6 weeks^24^. Body weights were monitored weekly for the first 6 weeks, then monitored every second day during the last 6 weeks of intervention to ensure accurate dosage for treatment. Blood samples were collected at baseline and biweekly (following a 4-6 h fast). After the completion of the 12 weeks, mice were anaesthetized with a ketamine and xylazine mixture (150 mg/kg ketamine and 10 mg/kg xylazine), exsanguinated by cardiac puncture, and perfused with PBS. Tissues of interest were removed and either snap-frozen in liquid nitrogen or placed in 10% formalin. The heart was carefully isolated and placed in 10% formalin and stored at 4 °C until processing.

**Cardiac Tissue Embedding and Sectioning:**

Hearts were fixed in 10% formalin for 48 hours at 4°C, while changing the formalin solution daily. Hearts were then incubated in sterile 20% sucrose in PBS for 72 hours at 4°C while changing the sucrose solution bi-daily. Before embedding, hearts were removed and dried gently of excess sucrose solution. A razor blade was used to cut transversely at roughly the central location of the heart. The upper chambers (containing the aortic sinus) was embedded in a tissue cassette with optimal cutting temperature medium, snap frozen with liquid nitrogen and stored at -80°C. The aortic sinus was sectioned using a cryostat (Leica CM 1850 cryostat) at 10 µm serial sections at -20 °C. Each slide contained (at least) 6 serial selections collected at 100 µm intervals, spanning at least 600 µm on a total of 10 slides and stored at -80 °C.

**Oil Red O Staining:**

Slides containing aortic sinus cross-sections were let equilibrate at room temperature for 2 min, following slides were fixed in 10% formalin in a Coplin jar for 10 min at room temperature. While fixation occurred, Oil Red O working solution (30 mL of 0.5% Oil Red O in isopropanol + 20 mL ddH_2_O) was prepared and let equilibrate for 10 min before being sequentially filtered (initial filtration with a coffee filter followed by filtration through a 0.45 µm syringe filter). Slides were washed once in 60% isopropanol and then stained in Oil Red O working solution for 10 min. Slides were washed twice with 60% isopropanol and then twice with PBS prior to hematoxylin counter staining. Slides were washed twice with ddH_2_O, once dry aqueous mounting media was added to the slide, and a 1 ½ cover slide was mounted. Slides were imaged at 20X magnification and tiled together with the EVOS FL auto 2. Atherosclerotic lesions were selected and quantified using Image J analysis software. Each data point is representative of the average plaque area quantified from 4 sections spanning 400 µm within the aortic sinus from one animal.

**Lesion Quantification and Analysis:**

Within the aortic sinus, lesions were stained with H&E and were selected and quantified using Image J software. Comparisons were made in tandem using a slide stained with Oil red O stained for increased confidence in plaque selection. Quantifications of plaque size were completed for each 10 µm section spanning at least 400 µm of the aortic sinus. The lesion size reported is the average lesion size from four sections spanning the same area within the aortic sinus for each experimental animal.

**Peritoneal macrophage isolation:**

Mice received an intraperitoneal injection of 1 mL sterile 3% thioglycolate medium four days prior to harvest to induce an inflammatory response for the recruitment of immune cells. At harvest, mice were euthanized by cervical dislocation and the abdominal skin was removed to expose the peritoneal cavity. Ten mL of ice-cold PBS was slowly injected within the peritoneal cavity through a 20G needle without removing the needle from the peritoneum, PBS and any free-floating peritoneal cells were slowly withdrawn back into the syringe. The 20G needle was removed and cells were slowly ejected into a sterile 50 mL conical tube and placed on ice until all isolations were completed. The cells were then centrifuged at 1000 RPM for 5 min at 4 °C, the supernatant was removed, and the cell pellet was gently re-suspended in 10 mL DMEM (supplemented with 10% FBS and 100 U/mL penicillin and streptomycin). Cells were seeded onto 1-3 100 mm cell culture dishes and let adhere for 16 h, then washed twice with PBS, gently scraped in complete DMEM and seeded onto 34.8 mm plates.

**Immunoblotting:**

Peritoneal cells were washed twice with ice-cold PBS and then cells were scrapped in 60-120 µL cell lysis buffer (50 mM Tris-HCl pH: 7.5, 150 mM, 1 mM EDTA, 0.5% Triton X-100, 0.5% NP-40, 100 µM Na_3_VO_4_, supplemented with protease inhibitor cocktail). All protein was quantified (via bicinchoninic acid assay - Pierce™), equalized, and 4X loading dye was added prior to denaturing the protein by boiling at 95 °C for 5 min. Protein samples were loaded and electrophoresed onto duplicate 8% SDS-PAGE gels (15-20 µg/well) for the determination of phosphorylated proteins along with their respective total amounts. Each gel was transferred onto PVDF membranes using the Trans-Blot Turbo system (25 V, 2.5 mps, for 18 min - Bio-Rad). Membranes were blocked in 5% BSA in TBST (w/v) with gentle rocking (Rocker 25 – Mendel) for 1 hour at room temperature, then incubated in the primary antibody of interest overnight at 4 °C. The following day membranes were washed 4 times in TBST at room temperature (5 min/wash), and membranes were incubated with anti-rabbit HRP conjugated secondary antibody for 1h at room temperature with gentle rocking and washed 4 more times in TBST. Clarity ECL substrate mix was added to membranes, which were imaged using the ImageQuant LAS4000 (GE) system. Quantification of total and relative luminescence for all proteins was done using Image J analysis software.

**Immunofluorescent Labeling:**

Slides containing aortic sinus cross-sections were let equilibrate to room temperature for 2 min and fixed in 2% PFA for 20 minutes. Following fixation, a hydrophobic marker (ImmunoPen - Millipore) was used to create barriers around all sections present on the slides. Slides were washed once in PBS then blocked/permeabilized with PBS containing 5% fatty-acid free BSA, 0.2% Triton-X100 and Tween-20. Samples were washed with PBS then co-stained with anti-mouse CD68-Alexa-647 (BioLegend) and p62-FITC (Novusbio) for 1 hour in antibody solution (2% BSA, 0.1% Triton X-100 and Tween-20 in PBS) at room temperature in a dehumidified chamber (shielded from light). Samples were washed twice with PBS then incubated with anti-FITC Alexa-488 conjugated secondary antibody (Invitrogen) for 1 hour at room temperature in a dehumidified chamber. Samples were washed with PBS then incubated in 300 nM DAPI (Thermo Fisher Scientific) for 5 min at room temperature shielded from light. Samples were washed twice with PBS, ProLong Gold antifade reagent (Invitrogen) was added to slides then mounted with 1 1/2 coverslip (Corning). Fluorescent microscopy images were taken using the EVOS FL Auto 2 Imaging System using a 20X magnification. All fluorescent images were processed and quantified using Image J analysis software.

**Plaque CD68 Composition:**

Firstly, we defined and selected lesions from immunofluorescent images using Image J software. Selected lesion areas were quantified for specific regional analysis. The lesions underwent thresholding to measure positive areas of fluorescence (where CD68 is expressed) within the plaque being defined as CD68+ area. The percent CD68+ area within the plaque was calculated as follows: ((CD68+ Lesion Area)/(Total Lesion Area))*100 = %CD68+ Plaque Area. Each reported value was the average of three sections within the aortic sinus of one animal, the same regions were quantified and compared for all animals.

**Lesion Autophagy:**

Firstly, we defined and selected lesions from immunofluorescent images using Image J software. Within selected regions, we quantified single-channel fluorescent intensity values corresponding to p62 expression. We analyzed the mean, median, and modal fluorescent intensities for the combined plaque within a section, only report the mean fluorescent intensity were reported. Each reported value was the average of three sections within the aortic sinus of one animal, the same regions were quantified and compared for all animals.

**Serum Lipid Determination:**

Mice were fasted for 4 hours prior to tail bleed (or cardiac puncture at experimental endpoint). Blood was left at room temperature for 30 min to allow to clot, then centrifuged at 3000 RPM for 10 min to allow for phase separation. The top opaque yellow phase (serum) was collected, aliquoted and stored at -20 °C. Serum was thawed and analyzed accordingly by the commercially available Infinity-cholesterol™ kit (Thermo Fisher Scientific) and the Triglyceride colorimetric assay kit (Cayman Chemical) as per manufacturer’s instructions.

**Serum Inflammatory Cytokine Analysis:**

Serum cytokines were quantified using the Meso Scale Discovery Pro-Inflammatory Panel 1 (mouse) kit (MSD, Gaithersburg, MD) assay using the manufacturer’s protocol. Serum monocyte chemoattractant protein 1 (MCP1) was determined by ELISA (R&D Systems), as per the manufacture’s instructions.

**Flow staining of immune populations:** Splenocytes were isolated by passing through nylon mesh strainers. Bone marrow cells were centrifuged from femurs and tibiae with cut epiphyses as previously described^26^. Blood was obtained via cardiac puncture and collected in EDTA-coated tubes. Erythrocytes were lysed in red blood cell lysis buffer (155 mM NH_4_Cl, 10 mM NaHCO_3_, 10 mM EDTA in H_2_O) with blood samples treated in 10 mL twice and splenocytes and bone marrow cells treated in 1 mL once. Cells were stained with Zombie Aqua (BioLegend) for 30 minutes on ice with anti-CD16/CD32 (unless included in surface staining) in PBS. Surface marker staining was performed in PBS + 0.5% BSA/2 mM EDTA/0.05% NaN3 (PBA-E) for 20 minutes on ice. Surface antibodies (from BioLegend unless otherwise specified) included: biotin-conjugated lineage cocktail (anti-B220, anti-Ter119, anti-CD11b, anti-Gr-1, anti-CD3ε; #133307), anti-Sca-1-PacificBlue (#108119), anti-c-Kit-PE/Cy7 (#105813), anti-CD34-eFluor660 (ThermoFisher #50-0341-82), anti-CD16/32-Brilliant Violet 711 (#101337), anti-CD115-PE (#135505), anti-Ly-6G-FITC (#127605), anti-Ly-6C-PE/Cy7 (#128017). Secondary stain with streptavidin-AlexaFluor 488 (BioLegend 405235) for 20 minutes on ice was done where necessary. Cells were fixed in 1-2% PFA in PBS for 15 minutes on ice and stored in PBA-E until acquired on an LSRFortessa cytometer (BD Biosciences). Analysis was performed using FlowJo VX (Treestar Inc.).

**Statistics:**

For atherosclerosis, immunofluorescent and biochemical analyses, all comparisons made between genotype and treatment, both within and between biological sex were made using 2-way ANOVA with a Tukey test for multiple comparisons (GraphPad Prism 7).
